# MARK3-mediated Slingshot-1 phosphorylation is essential for polarized lamellipodium formation

**DOI:** 10.1101/2023.10.15.562441

**Authors:** Toshiaki Mishima, Yusaku Ohta, Kazumasa Ohashi, Kensaku Mizuno

**Affiliations:** Laboratory of Molecular and Cellular Biology, Graduate School of Life Sciences, Tohoku University, Sendai, Miyagi 980-8578, Japan; Department of Chemistry, Graduate School of Science, Tohoku University, Sendai, Miyagi 980-8578, Japan

**Keywords:** Slingshot, cofilin, MARK3, 14-3-3 proteins, actin dynamics, lamellipodium, cell polarity, phosphorylation

## Abstract

Cofilin acts as a key regulator of actin cytoskeletal remodeling via stimulating actin filament disassembly. Cofilin is inactivated by Ser-3 phosphorylation and reactivated by cofilin-phosphatase Slingshot-1 (SSH1). SSH1 is activated upon binding to F-actin, and this activation is inhibited by its phosphorylation at Ser-937 and Ser-978 and the subsequent binding of 14-3-3 proteins. In this study, we identified MARK3 (also named Par-1a and C-TAK1) as a kinase responsible for Ser-937/Ser-978 phosphorylation of SSH1. MARK3-mediated phosphorylation promoted SSH1 binding to 14-3-3 proteins and suppressed its F-actin-assisted cofilin-phosphatase activity. When Jurkat cells were stimulated with SDF-1α, actin filaments formed multidirectional F-actin-rich lamellipodia around the cells in the initial stage, and thereafter, they were rearranged as a single polarized lamellipodium to the direction of cell migration. Upon SDF-1α stimulation, SSH1 was translocated into F-actin-rich lamellipodia, but its Ser-937/Ser-978 non-phosphorylatable mutant SSH1(2SA) was retained at the location of the original cortical F-actin. Knockdown of MARK3 or overexpression of SSH1(2SA), similar to SSH1 knockdown, impaired the conversion of multiple lamellipodia to a single polarized lamellipodium. These results indicate that MARK3-mediated Ser-937/Ser-978 phosphorylation is required for SSH1 liberation from F-actin and translocation to lamellipodia, and hence, facilitates the formation of a single polarized lamellipodium for directional cell migration. Our results suggest that the phosphorylation-dephosphorylation cycle of SSH1 is crucial for its localization to lamellipodia via promoting the dissociation-reassociation cycle of SSH1 to F-actin, and thereby the stimulus-induced lamellipodium formation to the direction of cell movement.

## Introduction

Actin filament dynamics and reorganization are essential for numerous cellular processes including morphological changes, polarity formation, and migration. Cofilin is a key regulator of actin cytoskeletal dynamics and reorganization via stimulating actin filament disassembly and severance (1, 2). Cofilin is inactivated by LIM-kinases (LIMKs)-mediated phosphorylation at Ser-3 (3, 4), and inactive Ser-3-phosphorylated cofilin (P-cofilin) is reactivated through dephosphorylation by the Slingshot (SSH) family of protein phosphatases, which consists of SSH1, SSH2, and SSH3 (5, 6). The spatial and temporal regulation of cofilin activity by these kinases and phosphatases plays crucial roles in cell morphological changes, polarity formation, and migration, as well as in many physiological and pathological processes, including embryogenesis, immune responses, neurogenesis, and cancer metastasis (1, 7).

When cells are stimulated with chemotactic factors, actin filaments are reorganized to produce multiple filamentous-actin (F-actin)-rich lamellipodial protrusions around the cell in the initial stage, and thereafter they are converted to a single polarized lamellipodium to the direction of cell migration; this protrusion is maintained at the leading-edge during cell migration (7–9). In the lamellipodium, there are polarized and dendritic arrays of actin filaments that are polymerized near the plasma membrane to push the membrane forward and depolymerized at the rear to replenish the actin monomers for further polymerization (10). It is conceivable that the spatiotemporally controlled change in cofilin activity through LIMK-mediated phosphorylation and SSH-mediated dephosphorylation is important for determining the rates of assembly and disassembly of actin filaments and the frequency of extension and retraction of lamellipodia, thereby regulating the production of multiple lamellipodia and their conversion to a single polarized lamellipodium after cell stimulation (7, 8). Depletion of SSH1 impairs the conversion of multidirectional lamellipodia to a single lamellipodium in stromal cell-derived factor-1α (SDF-1α)-stimulated Jurkat T cells (8), indicating that SSH1-mediated cofilin activation plays a crucial role in the formation of a single polarized lamellipodium for directional cell migration.

We previously showed that the cofilin-phosphatase activity of SSH1 is significantly increased by its association with F-actin (9, 11). Both the F-actin-binding activity and F-actin-assisted cofilin-phosphatase activity of SSH1 are suppressed by SSH1 phosphorylation at Ser-937 and Ser-978, and the subsequent binding of 14-3-3 proteins (9). 14-3-3 proteins bind to diverse proteins depending on the phosphorylation of their binding partners, and regulate their activity and localization (12, 13). We also showed that SSH1 accumulates in the F-actin-rich lamellipodia after cell stimulation with chemotactic factors (8, 9), suggesting that SSH1 is translocated to lamellipodia in response to cell stimulation and is locally activated via association with F-actin in lamellipodia, leading to cofilin-mediated actin remodeling in lamellipodia. Previous studies have shown that protein kinase D (PKD) family protein kinases phosphorylate SSH1 at Ser-937/Ser-978 and facilitate its binding to 14-3-3 proteins. This allows SSH1 sequestration in the cytoplasm and leads to the inhibition of cofilin activation and actin remodeling in lamellipodia, resulting in the impairment of directional cell migration (14, 15). These observations suggested that PKD-mediated SSH1 phosphorylation negatively regulates SSH1 activity and localization. However, the precise mechanism regulating SSH1 activity and localization and the functional consequences of Ser-937/Ser-978 phosphorylation of SSH1 remain to be fully elucidated.

In this study, we identified microtubule-associated regulatory kinase 3 (MARK3, also named Par-1a or C-TAK1) as the protein kinase responsible for Ser-937/Ser-978 phosphorylation of SSH1. MARK family kinases have been implicated in cell polarization, microtubule dynamics, and cell migration; however, their role in regulating actin filament dynamics remains largely unknown (16–18). We have shown that MARK3 phosphorylates SSH1 at Ser-937/Ser-978, promotes its binding to 14-3-3 proteins, and inhibits its F-actin-assisted cofilin phosphatase activity, similar to that in PKDs. However, contrary to the prediction that MARK3-mediated phosphorylation negatively regulates SSH1 activity and localization, we showed that a non-phosphorylatable mutant of SSH1 fails to localize to lamellipodia after cell stimulation and that depletion of MARK3 impairs the conversion of multiple lamellipodia to a single lamellipodium, similar to the phenotype of SSH1 depletion. These observations suggest that MARK3-mediated phosphorylation is required for SSH1 to be liberated from F-actin and translocated to the lamellipodia, leading to the formation of a single-polarized lamellipodium. We propose a model in which the cycle of phosphorylation and dephosphorylation of SSH1 is required for proper localization to the lamellipodia by stimulating the cycle of dissociation and reassociation of SSH1 to actin filaments.

## Results

### Screening of protein kinases responsible for Ser-978 phosphorylation of SSH1

The F-actin-binding and cofilin-phosphatase activities of SSH1 are inhibited by Ser-937/Ser-978 phosphorylation and subsequent binding of 14-3-3 proteins (9). To explore the regulatory mechanism of SSH1 activity and localization, we searched for the protein kinase(s) responsible for Ser-978 phosphorylation of SSH1 using an *E. coli* expression screening system (19) with the aid of an antibody specific to phospho-Ser-978 (pS-978) of SSH1. *E. coli* cells were transformed with the plasmid encoding glutathione S-transferase (GST)-fused SSH1 fragment (amino acids 869–1049) and then infected with a λ phage cDNA expression library prepared from mouse brain. Clones containing Ser-978-phosphorylated GST-SSH1(869–1049) were detected by immunoblotting using an anti-pS-978 antibody. Among 32 positive clones, 23 clones encoded protein kinases, including MARK3, calcium/calmodulin-dependent kinase (CaMK)-I, CaMK-II, protein kinase C-δ (PKC-δ), p21-activated kinase-1 (PAK1), CLK1 (also named STY), and phosphorylase kinase-γ. Of these protein kinases, we focused on MARK3 in this study, because the sequences surrounding Ser-937 and Ser-978 of SSH1 well accord with the consensus sequence motif of MARK-catalyzed phosphorylation sites, ΦxRxxS, where Φ is a hydrophobic residue (Fig. 1A), and MARK family kinases are known to promote the binding of the substrate proteins to 14-3-3 proteins (16–18).

**Figure 1.**
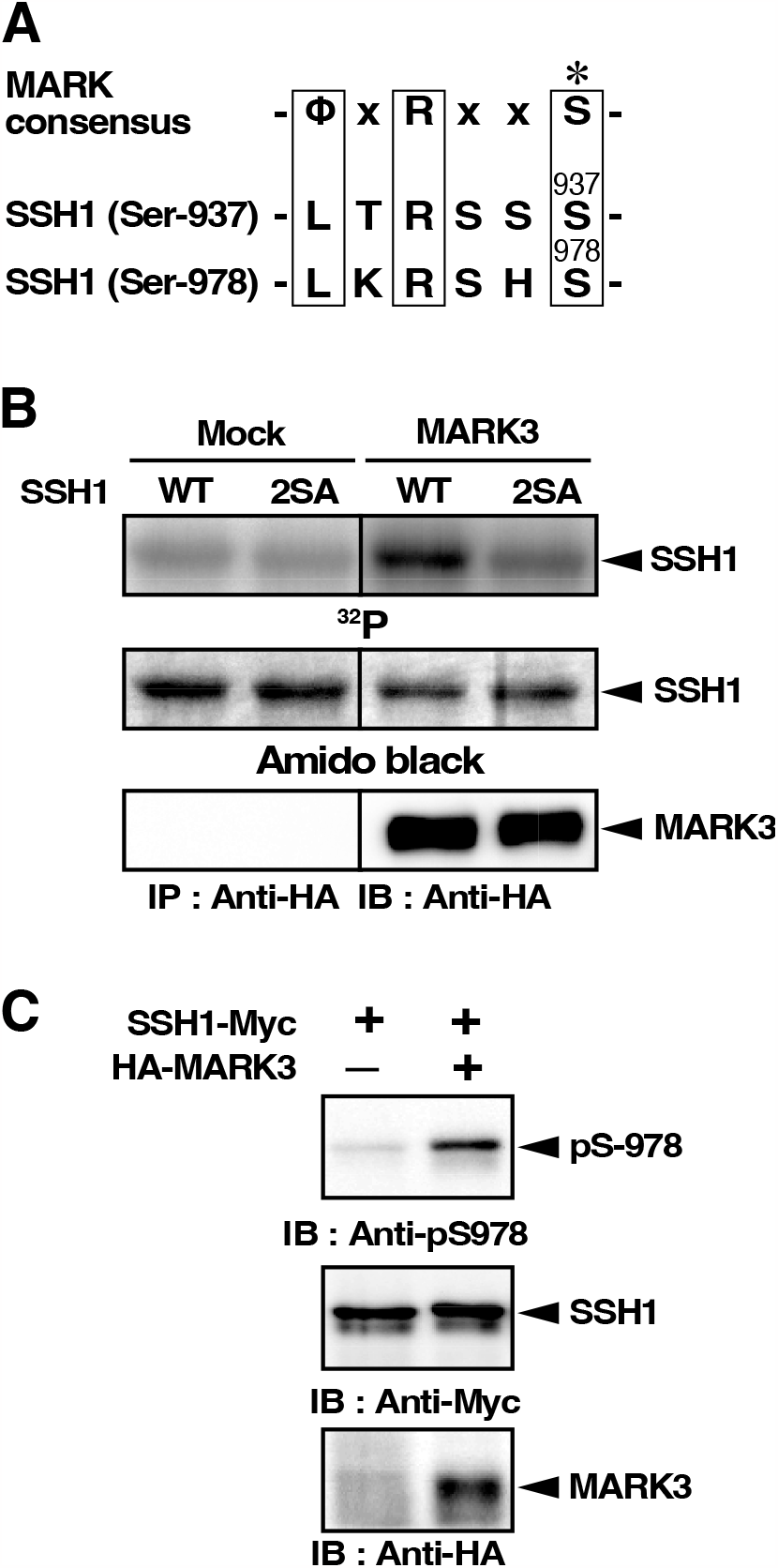
MARK3 phosphorylates SSH1 at Ser-978. (A) Alignment of the consensus sequence motif of the MARK3-catalyzed phosphorylation sites and sequences surrounding Ser-937 and Ser-978 of SSH1. *; phosphorylated serine, Φ; hydrophobic residue. (B) In vitro kinase assay. HA-tagged MARK3 was expressed in HEK293T cells, immunoprecipitated with anti-HA antibody, and subjected to in vitro kinase reaction, using [γ-^32^P]-ATP and recombinant [Myc-(His)_6_]-SSH1(WT or 2SA) as a substrate. Reaction mixtures were separated on SDS-PAGE and analyzed by autoradiography (^32^P), Amido black staining, and anti-HA immunoblotting. (C) MARK3 promotes Ser-978 phosphorylation of SSH1 in cultured cells. COS-7 cells were co-transfected with SSH1-Myc and HA-MARK3, and cell lysates were analyzed by immunoblotting with anti-pS-978, anti-Myc, and anti-HA antibodies.

### MARK3 phosphorylates SSH1 at Ser-978

To examine whether MARK3 actually phosphorylates Ser-937/Ser-978 of SSH1, HA-tagged MARK3 was expressed in HEK293T cells, immunoprecipitated with anti-HA antibody, and then subjected to *in vitro* kinase assays, using [γ-^32^P]-ATP and recombinant [Myc+(His)_6_]-tagged wild-type (WT) SSH1 or its 2SA mutant, in which Ser-937 and Ser-978 were replaced by alanine, as the substrates. [Myc+(His)_6_]-SSH1(WT) and its 2SA mutant were expressed in Sf9 cells and purified using nickel nitrilotriacetic acid (Ni-NTA)-agarose. *In vitro* kinase assays revealed that MARK3 effectively phosphorylated SSH1(WT), but not SSH1(2SA), indicating that MARK3 predominantly phosphorylated SSH1 at Ser-937/Ser-978 (Fig. 1B). To examine whether MARK3 phosphorylates Ser-978 of SSH1 in cultured cells, Myc-tagged SSH1 was co-transfected with HA-MARK3 in COS-7 cells and cell lysates were analyzed by immunoblotting with an anti-pS-978 antibody. The anti-pS-978-positive band was detected in MARK3-transfected cells, but not in MARK3-non-transfected cells, indicating that MARK3 promotes Ser-978 phosphorylation of SSH1 in cultured cells (Fig. 1C). Thus, MARK3 has the potential to phosphorylate Ser-978 of SSH1 both in cell-free assays and in cultured cells.

### MARK3 promotes SSH1 binding to 14-3-3 proteins

MARK kinases promote the binding of substrate proteins to 14-3-3 proteins and the sequences surrounding Ser-937 and Ser-978 correspond to the 14-3-3 binding motif RSxpS (pS, phospho-serine) (12, 13). Therefore, we examined whether the MARK3-catalyzed Ser-978 phosphorylation of SSH1 affects its binding to 14-3-3 proteins. [Myc+(His)_6_]-tagged-SSH1 (WT or 2SA) was expressed in Sf9 cells, purified using Ni-NTA-agarose, and then treated with or without MARK3 in a cell-free system. Reaction mixtures were then subjected to GST pull-down assays using GST-fused 14-3-3γ protein. The proteins bound to GST-14-3-3γ were analyzed by immunoblotting with anti-Myc and anti-pS-978 antibodies. This assay showed that treatment with MARK3 increased the level of Ser-978 phosphorylation of SSH1(WT) and its binding to GST-14-3-3γ (Fig. 2A). In contrast, SSH1(2SA) was neither phosphorylated nor pulled down by GST-14-3-3γ even after the treatment with MARK3 (Fig. 2A). These results suggest that MARK3-catalyzed phosphorylation of SSH1 at Ser-937/Ser-978 promotes its binding to 14-3-3γ. To further examine the role of MARK3 in SSH1 association with 14-3-3 proteins in cells, we analyzed the effect of MARK3 knockdown on their association. HEK293T cells were transfected with two independent MARK3-targeting short-hairpin RNAs (shRNAs) or control shRNA and cultured for 48 h. Cell lysates were analyzed by immunoprecipitation with an anti-SSH1 antibody, followed by immunoblotting with anti-SSH1 and anti-14-3-3 antibodies. Treatment with MARK3-targeting shRNAs reduced the expression of endogenous MARK3 protein in HEK293T cells (Fig. 2B). Co-immunoprecipitation assays with an anti-SSH1 antibody revealed that MARK3 knockdown significantly decreased the binding of endogenous 14-3-3 proteins to SSH1 (Fig. 2B). These results indicate that MARK3 plays a crucial role in the binding of SSH1 to 14-3-3 proteins by phosphorylating SSH1 at Ser-937/Ser-978.

**Figure 2.**
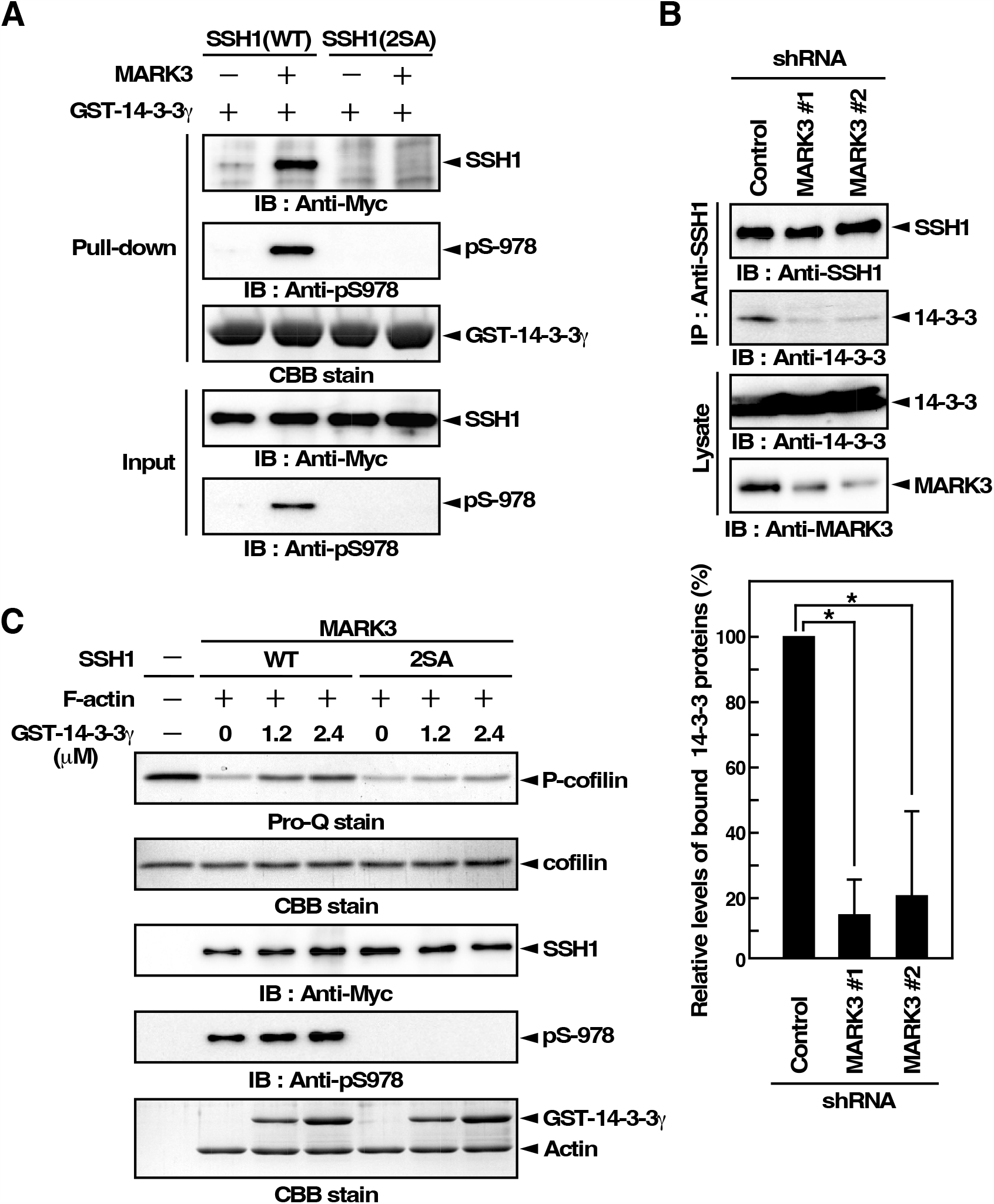
MARK3-mediated phosphorylation of SSH1 promotes its binding to 14-3-3 proteins and suppresses its cofilin-phosphatase activity. (A) MARK3-mediated Ser-937/Ser-978 phosphorylation of SSH1 promotes its binding to 14-3-3. HA-MARK3 was expressed in HEK293T cells, purified using anti-HA immunoprecipitation, and subjected to in vitro kinase reaction using recombinant [Myc+(His)_6_]-SSH1(WT or 2SA) as a substrate. After centrifugation, supernatants were subjected to in vitro pull-down assay with GST-14-3-3γ bound to glutathione-Sepharose, and the precipitates were analyzed by immunoblotting with anti-Myc and anti-pS-978 antibodies. GST-14-3-3γ was analyzed by CBB staining. The initial supernatants were directly subjected to SDS-PAGE and immunoblotted with anti-Myc and anti-pS-978 antibodies. (B) MARK3 knockdown reduces the interaction between SSH1 and 14-3-3 proteins in cells. HEK293T cells were transfected with the control or MARK3-targeting shRNAs. Endogenous MARK3 expression was analyzed by immunoblotting using an anti-MARK3 antibody. Cell lysates were immunoprecipitated with an anti-SSH1 antibody and the immunoprecipitates were analyzed by immunoblotting with anti-SSH1 and anti-14-3-3 antibodies. Quantitative data are shown as the means ± S.D. of three independent experiments. *, p<0.01. (C) MARK3-mediated Ser-937/Ser-978 phosphorylation and subsequent 14-3-3 binding suppress the cofilin-phosphatase activity of SSH1. [Myc+(His)_6_]-SSH1(WT or 2SA) was phosphorylated by HA-MARK3 as shown in (B). After centrifugation, the supernatants were incubated with indicated amount of GST-14-3-3γ, and then subjected to in vitro cofilin-phosphatase assay, using P-cofilin as a substrate, in the presence of F-actin. P-cofilin and total cofilin were detected using Pro-Q and CBB staining, respectively.

### MARK3-mediated SSH1 phosphorylation and its binding to 14-3-3 suppress F-actin-assisted cofilin-phosphatase activity of SSH1

The cofilin-phosphatase activity of SSH1 is markedly enhanced by its association with F-actin, which is inhibited by the binding of 14-3-3 proteins (9). To examine the role of MARK3 in SSH1 activity, we analyzed whether MARK3-mediated phosphorylation and subsequent 14-3-3γ binding affect the F-actin-assisted cofilin-phosphatase activity of SSH1. [Myc+(His)_6_]-SSH1(WT) or its 2SA mutant was expressed in Sf9 cells, purified using Ni-NTA-agarose, and treated with MARK3. MARK3-treated SSH1 (WT or 2SA) was subjected to an *in vitro* cofilin-phosphatase assay in the presence of F-actin and in the absence or presence of GST-14-3-3γ. The cofilin-phosphatase activity of SSH1(WT) was analyzed by measuring the decrease in the level of the substrate P-cofilin. The cofilin-phosphatase activity of MARK3-treated SSH1(WT) in the presence of F-actin was suppressed by the presence of 14-3-3γ in a dose-dependent manner (Fig. 2C). In contrast, 14-3-3γ had no apparent effect on the cofilin-phosphatase activity of MARK3-treated SSH1(2SA) in the presence of F-actin. These results suggest that MARK3-mediated Se-937/Ser-978 phosphorylation and subsequent binding of 14-3-3γ suppress F-actin-assisted cofilin-phosphatase activation of SSH1.

### Subcellular localization of SSH1 and its 2SA mutant

To examine the role of Ser-937/Ser-978 phosphorylation in cellular function of SSH1, we analyzed the subcellular localization of SSH1(WT) and its 2SA mutant. HeLa cells were transfected with [Myc+(His)_6_]-SSH1 or its 2SA mutant and co-stained with an anti-Myc antibody and rhodamine-labeled phalloidin to visualize F-actin. Both SSH1(WT) and SSH1(2SA) colocalized with F-actin, but their extent of colocalization differed (Fig. 3A). SSH1(2SA) almost entirely colocalized with F-actin throughout the cell. Compared to SSH1(2SA), SSH1(WT) colocalized with F-actin at a lower level, and some portion of SSH1(WT) was diffusely distributed in the cytoplasm (Fig. 3A). We also examined the localization of SSH1(WT) and SSH1(2SA) in Jurkat T cells. Jurkat cells were transfected with cyan fluorescent protein (CFP)-tagged SSH1(WT) or SSH1(2SA), and the localization of CFP-SSH1(WT or 2SA) and F-actin was visualized using CFP fluorescence and rhodamine-phalloidin staining, respectively. Confocal microscopic analysis of a cross-section of the vertical central region revealed that CFP-SSH1(2SA) mostly colocalized with cortical F-actin, but CFP-SSH1(WT) was localized in part to cortical F-actin, and the remainder was distributed in the cytoplasm (Fig. 3B). These observations suggest that SSH1(2SA), a Ser-937/Ser-978 non-phosphorylatable mutant of SSH1, colocalizes more tightly with F-actin than SSH1(WT), and that Ser-937/Ser-978 phosphorylation plays a role in dissociating SSH1 from F-actin structures in cells.

**Figure 3.**
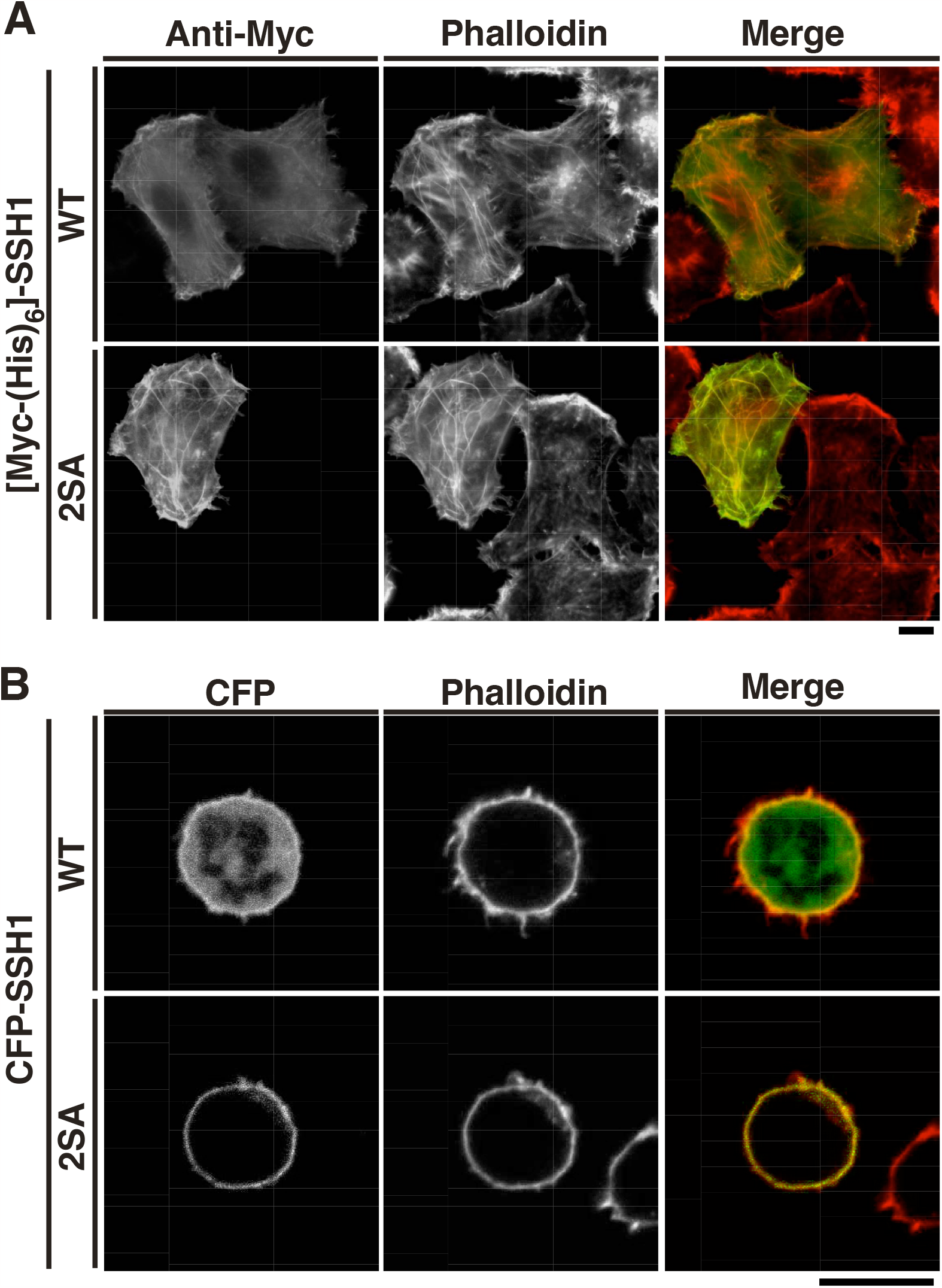
Subcellular localization of SSH1(WT) and SSH1(2SA) in HeLa and Jurkat cells. (A) HeLa cells transfected with [Myc+(His)_6_]-SSH1(WT or 2SA) were stained with anti-Myc antibody and rhodamine-phalloidin. Merged images of [Myc+(His)_6_]-SSH1 (WT or 2SA) (green) and rhodamine-phalloidin (red) are shown on the right. Scale bar, 10 μm. (B) Jurkat cells transfected with CFP-SSH1(WT or 2SA) were fixed and analyzed using CFP fluorescence and rhodamine-phalloidin staining. Confocal microscopic images of a cross-section of the vertical central region of the cell are shown. Merged images of CFP-SSH1(WT or 2SA) (green) and rhodamine-phalloidin (red) are shown on the right. Scale bar, 10 μm.

### Ser-937/Ser-978 phosphorylation is required for polarized lamellipodium formation in SDF-1α-stimulated Jurkat cells

We previously showed that knockdown of SSH1 suppresses the conversion of multidirectional lamellipodia to a single polarized lamellipodium in SDF-1α-stimulated Jurkat cells, suggesting the role of SSH1 for restricting the membrane protrusion to one direction (8). To examine the role of Ser-937/Ser-978 phosphorylation of SSH1 in polarized lamellipodium formation, we tested the effects of SSH1(WT or 2SA) overexpression on SDF-1α-induced changes in F-actin organization. Jurkat cells were transfected with CFP-SSH1(WT or 2SA) and then stimulated with SDF-1α. At 1, 5, and 20 min after stimulation, the cells were fixed and analyzed using CFP fluorescence and rhodamine-phalloidin staining. Three-dimensional projection images of phalloidin staining showed that cells expressing CFP-SSH1(WT) were round before stimulation, produced multiple lamellipodial protrusions at 1 min, and then became polarized to form a single lamellipodium in one direction at 5 and 20 min (Fig. 4A). In contrast, the majority of cells transfected with CFP-SSH1(2SA) produced multiple lamellipodial protrusions at 1 min and retained multiple lamellipodia even 5-20 min after SDF-1α stimulation (Fig. 4B). Thus, overexpression of SSH1(2SA) suppressed the conversion of multiple lamellipodia to a single polarized lamellipodium in SDF-1α-stimulated Jurkat cells, a phenotype similar to that induced by SSH1 knockdown.

**Figure 4.**
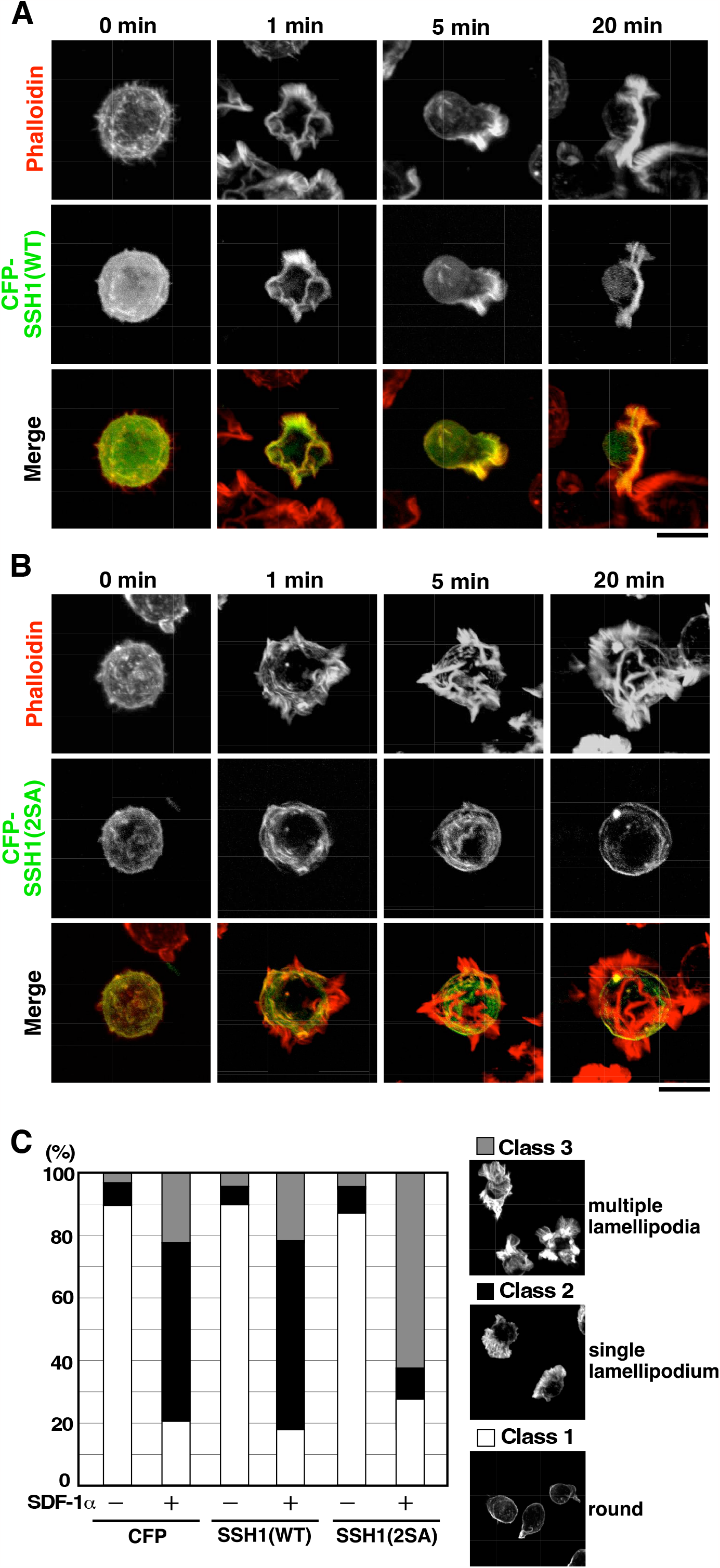
SSH1(2SA) does not translocate into lamellipodia and its overexpression impairs the conversion of multidirectional lamellipodia to a single polarized lamellipodium in SDF-1*α*-stimulated Jurkat cells. (A, B) Morphological changes and localization of SSH1(WT) (A) and SSH1(2SA) (B) in SDF-1α-stimulated Jurkat cells. Cells were unstimulated (0 min) or stimulated for 1, 5, and 20 min with 5 nM SDF-1α. The cells were fixed and analyzed using rhodamine-phalloidin staining and CFP fluorescence. Merged images of rhodamine-phalloidin (red) and CFP-SSH1(WT or 2SA) (green) are shown at the bottom. Scale bar, 10 μm. (C) Quantitative analysis of the cell morphology. Jurkat cells transfected with CFP, CFP-SSH1(WT), or CFP-SSH1(2SA) were stimulated with 5 nM SDF-1α for 15 min, fixed, and stained with rhodamine-phalloidin. Cells were categorized into three classes, as shown on the right: class 1 (round cells without a lamellipodium), class 2 (cells with a single lamellipodium), and class 3 (cells with multiple lamellipodia around the cells). The percentage of cells in each class is shown as the means of triplicate experiments (80-100 cells were counted in each experiment).

To quantify the effects of SSH1(WT) or SSH1(2SA) overexpression on polarized lamellipodium formation, cells were categorized into three classes, based on the F-actin organization and cell morphology: class 1 (round cells without a lamellipodium), class 2 (cells with a single lamellipodium), and class 3 (cells with multiple lamellipodia around the cells). Quantitative analyses showed that cells transfected with control CFP or CFP-SSH1(WT) preferentially exhibited cell morphologies categorized as class 2 at 15 min after SDF-1α stimulation (Fig. 4C). In contrast, cells transfected with CFP-SSH1(2SA) preferentially exhibited cell morphologies categorized as class 3 at 15 min after SDF-1α stimulation (Fig. 4C). These results suggest that Ser-937/Ser-978 phosphorylation of SSH1 is required for SDF-1α-induced polarized lamellipodium formation in Jurkat cells.

### Ser-937/Ser-978 phosphorylation is required for SDF-1α-induced SSH1 translocation into the lamellipodia

Confocal microscopic analyses of CFP fluorescence revealed that CFP-SSH1(WT) localized to the cortical F-actin and cytoplasm before SDF-1α stimulation (Fig. 3B and 4A) and accumulated in the F-actin-rich lamellipodial protrusions at 1–20 min after SDF-1α stimulation (Fig. 4A). Merged images show that SSH1(WT) mostly colocalized with F-actin in lamellipodial protrusions. SSH1(2SA) was more tightly colocalized with F-actin in the cell cortex of unstimulated cells (Fig. 3B and 4B). Notably, in contrast to SSH1(WT), SSH1(2SA) retained its original localization at cortical F-actin and did not translocate into lamellipodial protrusions after SDF-1α stimulation for 1–20 min (Fig. 4B). These results suggest that Ser-937/Ser-978 phosphorylation is required for SSH1 to translocate into the lamellipodial protrusions after SDF-1α stimulation. It is conceivable that SSH1 translocation into the lamellipodia is crucial for the conversion of multiple lamellipodia to a single polarized lamellipodium in SDF-1α-stimulated cells.

### MARK3 is required for SDF-1α-induced polarized lamellipodium formation

To examine the role of MARK3 in polarized lamellipodium formation, we analyzed the effect of MARK3 knockdown on SDF-1α-induced lamellipodium formation in Jurkat cells. Jurkat cells were transfected with control shRNA, SSH1-targeting shRNA, or two MARK3-targeting shRNAs, and cultured for 48 h. Immunoblot analyses showed that MARK3-targeting shRNAs effectively suppressed the expression of endogenous MARK3 protein in Jurkat cells (Fig. 5A). These cells were treated with SDF-1α for 15 min, and then fixed and stained with rhodamine-phalloidin, and their cell morphologies were analyzed as in Fig. 4C. Quantitative analyses showed that cells treated with MARK3 shRNAs preferentially exhibited cell morphology categorized into class 3 with multiple lamellipodia around the cells at 15 min after SDF-1α stimulation, whereas cells treated with control shRNA preferentially exhibited the morphology of class 2 with a single lamellipodium (Fig. 5B). As previously reported (8), cells treated with SSH1 shRNA preferentially exhibited cell morphology of class 3 after SDF-1α stimulation (Fig. 5B). These results indicate that similar to SSH1, MARK3 is required for the conversion of multiple lamellipodia to a single polarized lamellipodium in SDF-1α-stimulated Jurkat cells.

**Figure 5.**
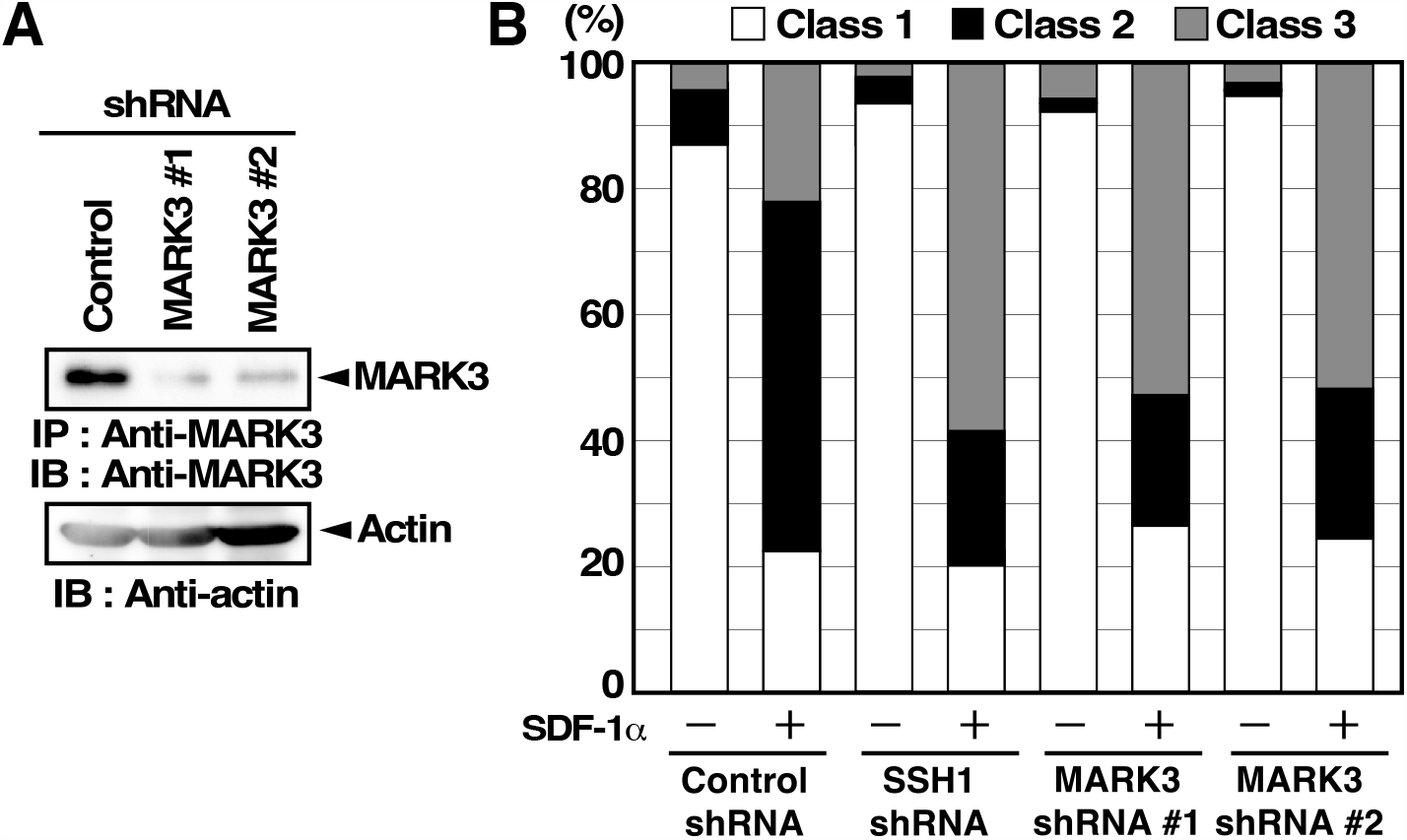
Knockdown of MARK3 impairs the conversion of multidirectional lamellipodia to a single polarized lamellipodium in SDF-1*α*-stimulated Jurkat cells. (A) Effects of MARK3-targeting shRNAs on the expression of MARK3. Jurkat cells were transfected with control or MARK3 shRNAs and cultured for 48 h. The cell lysates were immunoprecipitated and immunoblotted using an anti-MARK3 antibody. (B) Effects of MARK3 or SSH1 knockdown on SDF-1α-induced polarized lamellipodium formation. Jurkat cells were transfected with indicated shRNA plasmids, cultured for 48 h, and then stimulated with 5 nM SDF-1α for 15 min. The cells were then fixed and stained with rhodamine-phalloidin. Cells were categorized into three classes, as shown in Fig. 4C. The percentage of cells in each class is shown as the means of triplicate experiments (80-100 cells were counted in each experiment).

## Discussion

In this study, we identified MARK3 as a candidate protein kinase responsible for Ser-978 phosphorylation of SSH1. *In vitro* kinase assays revealed that MARK3 efficiently phosphorylated SSH1(WT), but not SSH1(2SA). In cultured cells, the level of Ser-978 phosphorylation of SSH1 was increased by coexpression of MARK3. These results indicate that MARK3 phosphorylates SSH1 at Ser-978. Since the sequences surrounding Ser-937 and Ser-978 of SSH1 are in accordance with the consensus sequence motif of the phosphorylation sites of MARK family kinases (16), MARK3 presumably phosphorylates SSH1 at both Ser-937 and Ser-978.

The MARK family kinases, consisting of MARK1/Par-1c, MARK2/Par-1b, MARK3/Par-1a, and MARK4/Par-1d in mammals, phosphorylate various proteins, thereby promoting their binding to 14-3-3 (16–18). We showed that MARK3 treatment promotes the binding of SSH1(WT), but not SSH1(2SA), to GST-14-3-3γ, and knockdown of MARK3 significantly suppressed the interaction between endogenous SSH1 and 14-3-3 proteins in cells, indicating that MARK3 promotes the association of SSH1 with 14-3-3 proteins, dependent on Ser-937/Ser-978 phosphorylation of SSH1. Furthermore, MARK3-mediated phosphorylation suppressed the F-actin-assisted cofilin-phosphatase activity of SSH1 in the presence of 14-3-3 proteins. Taken together, these results suggest that MARK3 plays a crucial role in regulating SSH1 association with 14-3-3 proteins and its cofilin-phosphatase activity. Because SSH1 binds to F-actin through the C-terminal serine-rich region (amino acids 935-985), including Ser-937 and Ser-978 (20), it is conceivable that 14-3-3 proteins inhibit F-actin-assisted activation of SSH1 by blocking the binding of SSH1 to F-actin.

Microscopic analyses revealed that SSH1(2SA) almost completely co-localizes with F-actin, whereas SSH1(WT) localizes both on F-actin and in the cytoplasm, suggesting that SSH1 tightly binds to F-actin in its Ser-937/Ser-978 non-phosphorylated form but it is dissociated from F-actin when it is phosphorylated at Ser-937/Ser-978 and subsequently associated with 14-3-3 proteins. Thus, MARK3-mediated phosphorylation of Ser-937/Ser-978 seems to play an important role in determining the subcellular localization of SSH1. As reported previously (8), SSH1(WT) was accumulated in the lamellipodial protrusions after SDF-1α stimulation. In contrast, SSH1(2SA) did not translocate into the lamellipodial protrusions and retained in the cortical F-actin even after SDF-1α stimulation. The marked difference in the localization of SSH1(2SA) and SSH1(WT) strongly suggests that Ser-937/Ser-978 phosphorylation, followed by 14-3-3 binding, is required for SSH1 liberation from actin filaments and its translocation into the lamellipodial protrusions (Fig. 6).

**Figure 6.**
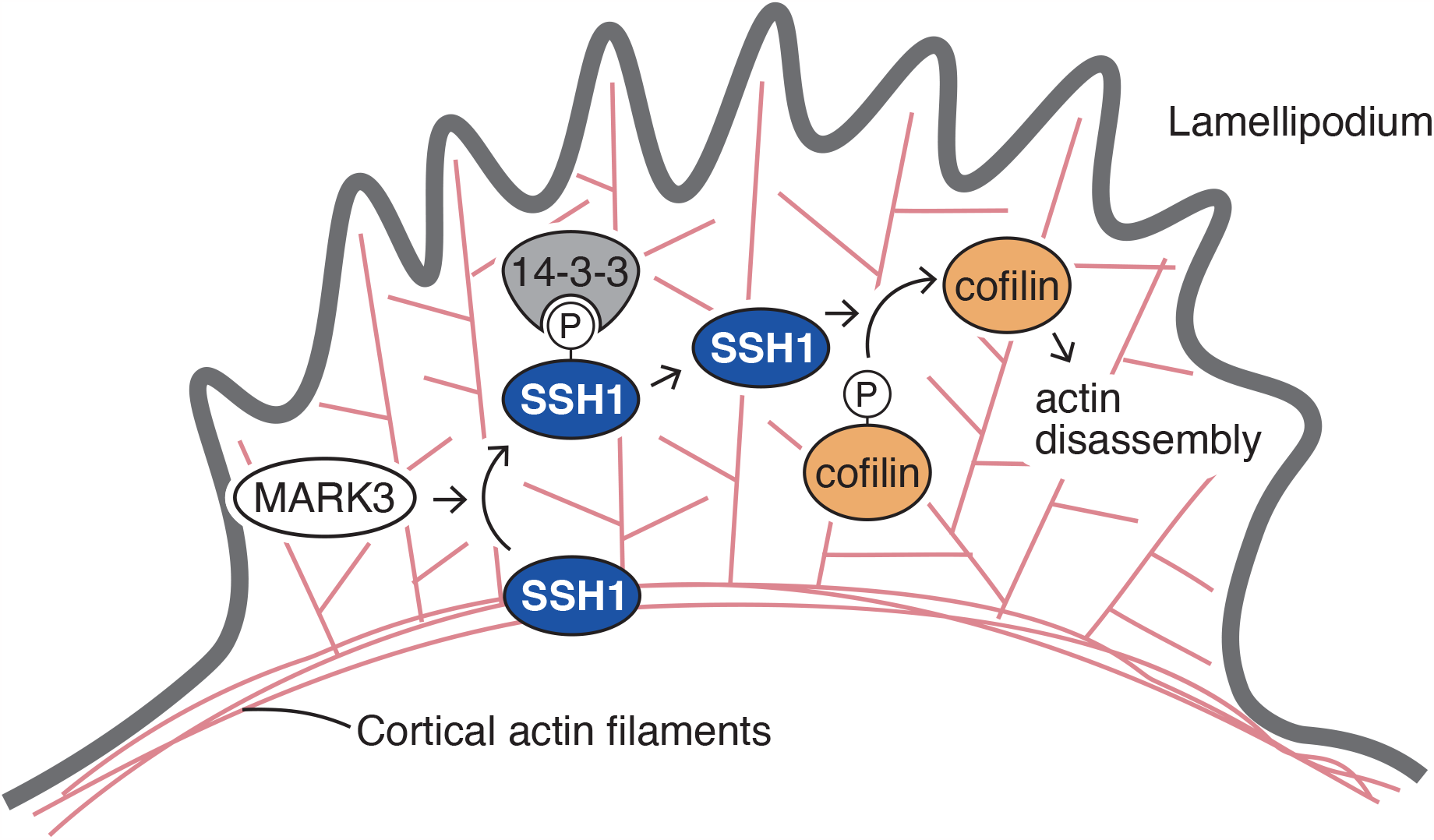
A model of phospho-regulation of SSH1 localization in SDF-1α-induced polarized lamellipodium formation in Jurkat cells. In response to SDF-1α stimulation, SSH1 is translocated from the cortical actin filaments to the inside of the lamellipodium. MARK3-mediated Ser-937/Ser-978 phosphorylation of SSH1 and subsequent binding to 14-3-3 enable SSH1 liberation from cortical F-actin and translocation into the lamellipodia, thereby supporting the formation of a single polarized lamellipodium in the direction of cell movement.

Stimulation of Jurkat cells with SDF-1α induces the formation of multiple lamellipodia around the cell in the initial stage of cell response, and then multiple lamellipodia are rearranged to a single polarized lamellipodium in the later stage; this single lamellipodium is maintained at the leading edge of migrating cells. We previously showed that knockdown of SSH1 suppresses this rearrangement in SDF-1α-stimulated Jurkat cells (8), indicating that SSH1-mediated cofilin activation is required for restricting the lamellipodial protrusion to one direction by abolishing the other lamellipodia. Because SSH1 is required to restrict lamellipodial protrusion in one direction, it is predicted that the failure of SSH1 to translocate to the lamellipodia impairs the conversion of multiple lamellipodia to a single polarized lamellipodium. Depletion of MARK3 or overexpression of SSH1(2SA) suppressed the formation of a single-polarized lamellipodium in SDF-1α-stimulated Jurkat cells, indicating that MARK3-mediated SSH1 phosphorylation at Ser-937/Ser-978 is required for this process. It is likely that phosphorylated SSH1 binds to 14-3-3 proteins and is translocated into the lamellipodia; thereafter, it is dephosphorylated and released from 14-3-3 proteins by unknown mechanisms and then activated by association with F-actin, which in turn promotes cofilin activation and F-actin disassembly in lamellipodia, leading to the conversion of multiple lamellipodia to a single polarized lamellipodium (Fig. 6). Although MARK3-mediated SSH1 phosphorylation negatively regulates the cofilin-phosphatase activity of SSH1, MARK3 depletion resulted in a phenotype similar to that of SSH1 depletion. This is probably because MARK3 depletion suppresses SSH1 release from cortical F-actin and its translocation into lamellipodia. Thus, it is conceivable that the cycle of phosphorylation and dephosphorylation of SSH1 is involved in SSH1 translocation into lamellipodia, cofilin activation in lamellipodia, and polarized lamellipodium formation. Further studies are needed to understand the mechanism by which SSH1 is released from 14-3-3 proteins and the types of protein phosphatases involved in Ser-937/Ser-978 dephosphorylation in lamellipodia.

Previous studies have shown that PKD family protein kinases phosphorylate SSH1 at Ser-937 and Ser-978, which leads to its association with 14-3-3 proteins and the inhibition of its cofilin-phosphatase activity (14, 15). Based on the observation that the expression of active PKD1 suppressed and the depletion of PKD1 promoted directional cell migration, they proposed a model in which PKD-mediated SSH1 phosphorylation negatively regulates directional cell migration by inhibiting SSH1-mediated cofilin activation in lamellipodia (14, 15). In contrast, we showed that the depletion of MARK3 inhibits the rearrangement of multiple lamellipodia around the cell to a single polarized lamellipodium in one direction, which is an essential step for directional cell migration. We propose a model in which the phosphorylation-dephosphorylation cycle of SSH1 is involved in its proper localization to lamellipodia by promoting the dissociation-reassociation cycle of SSH1 to F-actin, thereby promoting the formation of a single polarized lamellipodium. As for the difference between these studies, PKD promotes cofilin phosphorylation and inactivation through activation of the PAK-LIMK pathway, in addition to direct phosphorylation and inactivation of SSH1 (21). It was also shown that PKD phosphorylates SSH1 at Ser-402 in the phosphatase catalytic domain, which causes inactivation of its cofilin-phosphatase activity (22). These characteristics of PKD may explain why PKD depletion results in phenotypes distinct from those induced by MARK3 depletion.

MARK family kinases are mammalian orthologs of Par-1 (partitioning defective-1) that was originally identified in *Caenorhabditis elegans* as a key regulator of cell polarity formation (23). Par-1 is asymmetrically localized in the posterior cortex and plays an essential role in establishing the anterior/posterior axis in *C. elegans* embryos and *Drosophila* oocytes (24–26). Par-1 restricts the localization of the anterior Par complex, consisting of Par-3, Par-6, and aPKC, to the anterior region by phosphorylating Par-3, whereas aPKC phosphorylates Par-1 and restricts its localization to the posterior region (26). In mammals, MARK kinases were first identified by their ability to phosphorylate microtubule-associated proteins (MAPs) (27). MARK kinases are required for apical-basolateral polarity formation in epithelial cells, axon/dendrite specification, dendritic spine maturation in neurons, and directional migration of various cell types, such as astrocytes, fibroblasts, cortical neurons, and cancer cells (16–18). For cell migration, Par-1/MARK kinases play a role in establishing the front-rear orientation of migrating cells, probably by supporting the localization of the Par-3/Par-6/aPKC complex at the front and by regulating polarized microtubule network formation via the phosphorylation of Par-3 and MAPs, respectively (17, 28). In *Drosophila* border cell migration during ovary development, Par-1 mutations result in misoriented protrusions around the cell, leading to defects in directional cell migration, indicating that Par-1 is required for destabilizing protrusions at the rear and forming a polarized protrusion in the direction of cell migration (29). These phenotypes of border cells were closely related to those observed upon MARK3 depletion in Jurkat cells shown in this study. Thus, the function of PAR-1/MARK kinases in producing a single polarized lamellipodium for directional cell migration appears to be conserved in *Drosophila* and human cells.

Actin cytoskeletal reorganization is essential for the front-rear polarity formation in migrating cells. A recent study showed that MARK2 is required for directional cell migration by promoting phosphorylation of the myosin II regulatory light chain and myosin phosphatase targeting subunit MYPT1, which induces actomyosin contractility (30). Here, we show that MARK3-mediated SSH1 phosphorylation is required for the proper localization of SSH1 in lamellipodia, which promotes actin filament dynamics in lamellipodia and polarized lamellipodium formation. The identification of SSH1 as a downstream effector of the MARK signaling pathway will provide new insights into the molecular mechanisms of MARK kinase-mediated cell polarity formation and directional cell migration. However, the question of how a single lamellipodium is selectively produced at the front of the migrating cell from the initially formed multidirectional lamellipodia remains unanswered and warrants further investigation.

## Experimental procedures

### Plasmids and antibodies

Expression plasmids coding for C-terminally [Myc+(His)_6_]- or N-terminally CFP-tagged SSH1(WT) and SSH1(2SA) and GST-14-3-3γ were constructed, as described previously (8, 9). The cDNA plasmid coding for 14-3-3γ was provided by T. Ichimura and T. Isobe (Tokyo Metropolitan University). Expression plasmids coding for N-terminally HA-tagged MARK3 was constructed by inserting the cDNA of MARK3 (amino acids 52-798) into FPC-HA vector. For protein expression in baculovirus system, cDNA inserts were subcloned into pFastBac1 vector (Invitrogen) as described previously (31). For shRNA, oligonucleotides were annealed and subcloned into pSUPER vector plasmids (provided by R. Agami, Netherland Cancer Institute) as described previously (32). The 19-base targeting sequences used in this study were as follows: 5’-CGCGGCACTCTAGAGCAAA-3’ (MARK3 #1), 5’-GCCGAGGCTCCACTAATCT-3’ (MARK3 #2), 5’-TCGTCACCCAAGAAAGATA-3’ (SSH1), and 5’-TCTTCCCCCAAGAAAGATA-3’ (as a control). Antibodies against Myc (9E10; Roche), 14-3-3 (H-8; Santa Cruz Biotechnology), MARK3 (G-7; Santa Cruz Biotechnology), and β-actin (AC15, Millipore-Sigma) were purchased commercially. Antibodies against SSH1, cofilin, and pS-978 were prepared as described previously (9).

### Cell culture and transfection

HEK293T, COS-7 and HeLa cells were cultured in Dulbecco’s modified Eagle’s medium (DMEM) containing 10% fetal calf serum (FCS). Cells were transfected with plasmid using FuGENE6 Tansfection Reagent (Roche) or Lipofectamine (Invitrogen) according to the manufacturer’s instructions. Jurkat cells were maintained in Roswell Park Memorial Institute (RPMI) 1640 medium supplemented with 9% FCS. For transfection into Jurkat cells, ∼1 × 10^7^ cells were mixed with plasmids in 400 μl of electroporation medium (RPIM 1640 containing 20% FCS and 25 mM Hepes, pH 7.4) and electroporated at 280 V and 725 μF using a Gene Pulser II (Bio-Rad Laboratories). After electroporation, the cells were cultured for 36–60 h in RPMI 1640 medium supplemented with 10% FCS.

### Expression cloning

To identify the kinase(s) responsible for Ser-978 phosphorylation of SSH1, we performed expression cloning in *E. coli*, as described previously (19). The cDNA encoding GST-SSH1 (869–1049) was constructed by inserting the cDNA for human SSH1 (869–1049) into the *Bam*HI and *Not*I sites of pGEX-4T (GE Healthcare). *E. coli* cells were transformed with the plasmid for GST-SSH1 (869–1049) and then infected with a λ phage cDNA library from neonatal mouse brain (λ Uni-ZAP XR, Stratagene). ExAssist interference-resistant helper phages were coinfected with λ phage library to induce *in vivo* excision of cDNA into pBluescript phagemid in *E. coli* cells. Expression of GST-SSH1(869–1049) and library-oriented proteins was induced by overlaying to polyvinylidene fluoride (PVDF) membrane, which had presoaked in 20 mM isopropyl β-D-1-thiogalactopyranoside (IPTG), and incubating on the plates at 37? for 4 h. The membranes were washed with Tween-phosphate-buffered saline (PBS) (1.37 mM NaCl, 27 mM KCl, 81 mM Na_2_HPO_4_, 15 mM KH_2_PO_4_, 1% Tween 20), and blocked with Tween-PBS containing 5% skim milk for 1 h. The membranes were then immunoblotted with anti-pS-978 antibody and visualized by horseradish peroxidase (HRP)-labeled secondary antibody and Chemi-Lumi One L (Nacalai tesque, Japan).

### In vitro kinase assay

Recombinant [Myc+(His)_6_]-SSH1(WT or 2SA) was expressed in Sf9 cells and purified by using Ni-NTA-agarose, as described previously (31). HA-tagged MARK3 was expressed in HEK293T cells and purified by immunoprecipitation with anti-HA antibody. Immunoprecipitates were washed with kinase buffer (25 mM Tris-HCl, pH 7.5, 150 mM NaCl, 5% glycerol, 1 mM dithiothreitol, 20 mM NaF, 1 mM Na_3_VO_4_, 1 mM phenylmethylsulfonyl fluoride, 10 μg/ml leupeptin, 1 mM MgCl_2_, 1 mM MnCl_2_) and incubated with recombinant [Myc+(His)_6_]-SSH1 (WT or 2SA), 50 μM ATP, and 185 kBq [γ-^32^P]ATP(3000 Ci/mmol, Perkin-Elmer) in kinase buffer at 30? for 1 h. The reaction mixtures were separated by SDS-PAGE and analyzed by autoradiography to measure ^32^P-labeled SSH1.

### GST pull-down assay

HA-MARK3 was expressed in HEK293T cells, purified by immunoprecipitation with anti-HA antibody. Immunoprecipitates were washed with kinase buffer and incubated in kinase buffer containing 50 μM ATP with recombinant [Myc+(His)_6_]-SSH1 at 30? for 1 h. After kinase reaction, recombinant GST-14-3-3γ was added to reaction mixtures and incubated at 4? for 1 h. After centrifugation, the supernatants were incubated with glutathione-Sepharose (GE Healthcare) at 4? for 1 h. Then the beads were washed with lysis buffer (25 mM Tris-HCl, pH 7.5, 150 mM NaCl, 1% Nonidet P-40, 5% glycerol, 1 mM dithiothreitol, 20 mM NaF, 1 mM Na_3_VO_4_, 1 mM phenylmethylsulfonyl fluoride, 10 μg/ml leupeptin, 1 mM MgCl_2_) and subjected to SDS-PAGE and immunoblot analysis.

### In vitro cofilin-phosphatase assay

Cofilin-phosphatase activity of SSH1 was analyzed as described previously (8). Cofilin-(His)_6_ was expressed in Sf9 cells, purified with Ni-NTA agarose. Recombinant [Myc+(His)_6_]-SSH1 was purified and phosphorylated by HA-MARK3, and then incubated with 0–2.4 μM GST-14-3-3γ for 30 min at 30°C, and then incubated with P-cofilin-(His)_6_ in phophatase buffer (25 mM Tris-HCl, pH 7.5, 150 mM NaCl, 1% Nonidet P-40, 5% glycerol, 1 mM dithiothreitol, 10 μg/ml leupeptin, 1 mM MgCl_2_) containing 250 μg/ml F-actin for 5 min at 30?. Reaction mixtures were subjected to SDS-PAGE. P-cofilin and total cofilin were analyzed by Pro-Q Diamond phosphoprotein gel stain kit (Invitrogen) and Coomassie brilliant blue (CBB) staining, respectively.

### Cell staining

HeLa cells were fixed with 4% formaldehyde in PBS for 30 min and permeabilized with 1% Triton X-100 in PBS containing 2% FCS for 5 min. After blocking with PBS containing 2% FCS for 1 h, cells were stained with anti-Myc antibody followed by staining with fluorescein-isothiocyanate (FITC)-labeled anti-mouse IgG antibody (Chemicon). Jurkat cells were suspended in RPMI 1640 medium containing 25 mM Hepes (pH 7.4) and 0.2% bovine serum albumin and incubated for 30 min at 37?. Cells were stimulated with 5 nM SDF-1α for 0–20 min, fixed with formalin for 15 min, permeabilized with 1% Triton X-100 in PBS containing 2% FCS for 5 min, blocked with PBS containing 2% FCS for 30 min, and then analyzed by CFP fluorescence. F-actin was stained with rhodamine-conjugated phalloidin (Molecular Probes).

## Footnotes

## Acknowledgments

We thank Dr. R. Agami (Netherland Cancer Institute) for providing pSUPER vector and Drs. T. Ichimura and T. Isobe (Tokyo Metropolitan University) for providing cDNA plasmids encoding 14-3-3.

## Author contributions

T. M., Y. O., and K. O. resources; T. M. and Y. O. data curation; T. M. and Y. O. formal analysis; T. M., Y. O., and K. M. validation; T. M. and Y. O. investigation; T. M. visualization; T. M., Y. O., and K. O. methodology; T. M. and K. M. conceptualization; K. O. and K. M. supervision; K. O. and K. M. funding acquisition; T. M. and K. M. writing-original draft; K. M. project administration; K. M. writing-review and editing.

## Funding and additional information

This work was supported by Grants-in aid for Scientific Research 20H03248 and 22H05618 (to K. O.) and 16H00749 and 18K19280 (to K. M.) from the Japan Society for the Promotion of Science (JSPS), and The Uehara Memorial Foundation (to K. O.).

## Conflict of interest

The authors declare that they have no conflicts of interest with the contents of this article.

### Abbreviations

CFP: cyan fluorescence protein
F-actin: filamentous actin
FCS: fetal calf serum
GST: glutathione S-transferase
IPTG: isopropyl β-D-1-thiogalactopyranoside
LIMK: LIM-kinase
MARK: microtubule affinity-regulating kinase 3
Ni-NTA: nickel nitrilotriacetic acid
PAK: p21-activated kinase
PAR-1: partitioning defective-1
PBS: phosphate-buffered saline
P-cofilin: Ser-3-phosphorylated cofilin
PKC: protein kinase C
PKD: protein kinase D
pS-978: phospho-serine 978
SDF-1α: stromal cell-derived factor 1α
shRNA: short hairpin RNA
SSH1: Slingshot 1
SSH1(2SA): SSH1 mutant in which Ser-937 and Ser-978 are replaced by alanine
WT: wild-type

